# Haplotype-resolved genome assembly of the tetraploid potato cultivar Désirée

**DOI:** 10.1101/2025.01.14.631659

**Authors:** Tim Godec, Sebastian Beier, Natalia Yaneth Rodriguez-Granados, Rashmi Sasidharan, Lamis Abdelhakim, Markus Teige, Björn Usadel, Kristina Gruden, Marko Petek

## Abstract

Cultivar Désirée is an important model for potato functional genomics studies to assist breeding strategies. Here, we present a haplotype-resolved genome assembly of Désirée, achieved by assembling PacBio HiFi reads and Hi-C scaffolding, resulting in a high-contiguity chromosome-level assembly. We implemented a comprehensive annotation pipeline incorporating gene models and functional annotations from the *Solanum tuberosum* Phureja DM reference genome alongside RNA-seq reads to provide high-quality gene and transcript annotations. Additionally, we provide a genome-wide DNA methylation profile using Oxford Nanopore reads, enabling insights into potato epigenetics. The assembled genome, annotations, methylation and expression data are visualised in a publicly accessible genome browser (https://desiree.nib.si), providing a valuable resource for the potato research community.

## Background & Summary

Potato (*Solanum tuberosum*) is one of the most important and widely cultivated crops worldwide, with a significant role in global food security and agricultural research. Despite its significance, many studies still rely on the genome of the double monoploid (DM) clone of group Phureja DM1–3 516 R44^1,2^ which lacks a substantial portion of the gene repertoire and variability found in cultivated tetraploid potato varieties.

The potato cultivar Désirée is a red-skinned late-season potato variety, originally bred in the Netherlands in 1962 by crossing parent cultivars Urgenta and Depesche (Potato Pedigree Database)^3^. It is still cultivated due to its favourable agronomic traits, such as predictable yields and high tolerance to drought and some pathogens^4^. It has also been used in breeding programs, yet a genome assembly for the Désirée cultivar has not been available. In research, it has been propagated in tissue cultures, and used for genetic manipulation including gene overexpression^5^, gene silencing^6^, and Crispr-Cas gene editing^7^.

Although haplotype-resolved genome assemblies are becoming common in diploid organisms, the high heterozygosity rate, extensive repeat content, and the autopolyploid nature of cultivated potatoes still present significant challenges for generating high-quality haplotype-resolved assemblies. Currently, five haplotype-resolved genomes of autotetraploid potato cultivars are publicly available^8–12^ as well as several phased diploid genomes^13–15^. The recently published haplotype-resolved tetraploid potato assemblies rely on labour-intensive techniques such as single-pollen sequencing^10^ or the use of parental and crossing material^11^, which may not always be available.

Adding to existing publicly available genomes, we provide a reference quality (CRAQ overall AQI of 97.5) haplotype-resolved genome assembly of the tetraploid cultivar Désirée, assembled using solely PacBio HiFi and Illumina Hi-C data. Our assembly is accompanied by a comprehensive structural and functional gene annotation reaching 99.4 % BUSCO completeness for Solanaceae, accompanied by orthology to DM genes. For the potato research community, we provide an online resource featuring a genome browser and downloadable genomic assembly and annotation files, providing a valuable tool for studies involving allele-specific expression or promoter analysis.

## Methods

### Sample preparation and sequencing

Leaves from 4-week old *S. tuberosum* cv. Désirée plants were collected and flash-frozen. High molecular weight genomic DNA (HMW gDNA) used for PacBio HiFi, Illumina and Oxford Nanopore Technologies (ONT) sequencing was extracted from the leaf tissues using a modified CTAB method^16^. The concentration and quality of the extracted DNA were assessed using a NanoDrop spectrophotometer.

#### PacBio HiFi

HMW gDNA was sent to National Genomics Infrastructure (NGI) Sweden for library preparation and sequencing on the PacBio Sequel II platform. We obtained 79.4 Gbp of raw data, consisting of 4.1 million reads.

#### Illumina Hi-C

Leaves from 4-week old *S. tuberosum* cv. Désirée plants were collected, flash-frozen in liquid nitrogen and ground using mortar and pestle. Hi-C library prep using the Omni-C kit (Dovetail Genomics) and sequencing were performed on an Illumina NovaSeq 6000 platform by NGI Sweden. Sequencing generated 2018.4 million paired-end (2 × 150 bp) reads.

#### ONT

The HMW gDNA was used for ONT DNA library prep using the SQK-LSK110 kit and sequenced on a MinION using the FLO-MIN106 flow cell. Reads were basecalled using Dorado (v0.7.2) with the model dna_r9.4.1_e8_sup@v3.3 which generated 5.8 Gbp. The reads with methylation-related tags were converted to bedMethyl format using modkit (v0.4.1).

#### Illumina short reads

Illumina short-read library was constructed from the HMW gDNA and sequenced on Illumina NextSeq 2000 by ELIXIR Slovenia node to generate 150 bp paired-end reads. The short-read sequencing generated approximately 138 Gbp of raw data, consisting of 460.1 million paired-end (2 × 150 bp) reads.

### Genome size and heterozygosity estimation

The genome characteristics of *S. tuberosum* cv. Désirée, including genome size, heterozygosity, and repeat content, were estimated using Illumina short-read data and a k-mer based approach. A 21-mer frequency distribution was generated with Jellyfish (v2.2.10), and the genome’s key features were inferred using GenomeScope2 (v2.0). The haploid genome size was estimated at 669.6 Mbp, with a heterozygosity rate estimated at 3.8–5.7%.

### *De novo* genome assembly, Hi-C scaffolding and quality assessment

PacBio HiFi and Illumina Hi-C reads were initially assembled into four sets of haplotype-resolved contigs using Hifiasm (v0.19.8-r603)^17–19^. Hifiasm primary unitigs were searched against DM genome assembly with blastn (v2.5.0)^20^ and best matches were visualised on Graphical Fragment Assembly with Bandage (v0.8.1, Fig. 1a)^21^. We performed quality control of the contigs using Merqury (v1.3, Fig. 1b)^22^ k-mer spectra and BUSCO completeness scores (v5.4.7, solanales_odb10 dataset)^23^. The length of haplotype draft assemblies ranged from 761.6 Mbp to 888.4 Mbp with contig N50 sizes ranging from 7.0 Mbp to 13.7 Mbp (Table 1).

**Fig. 1.**
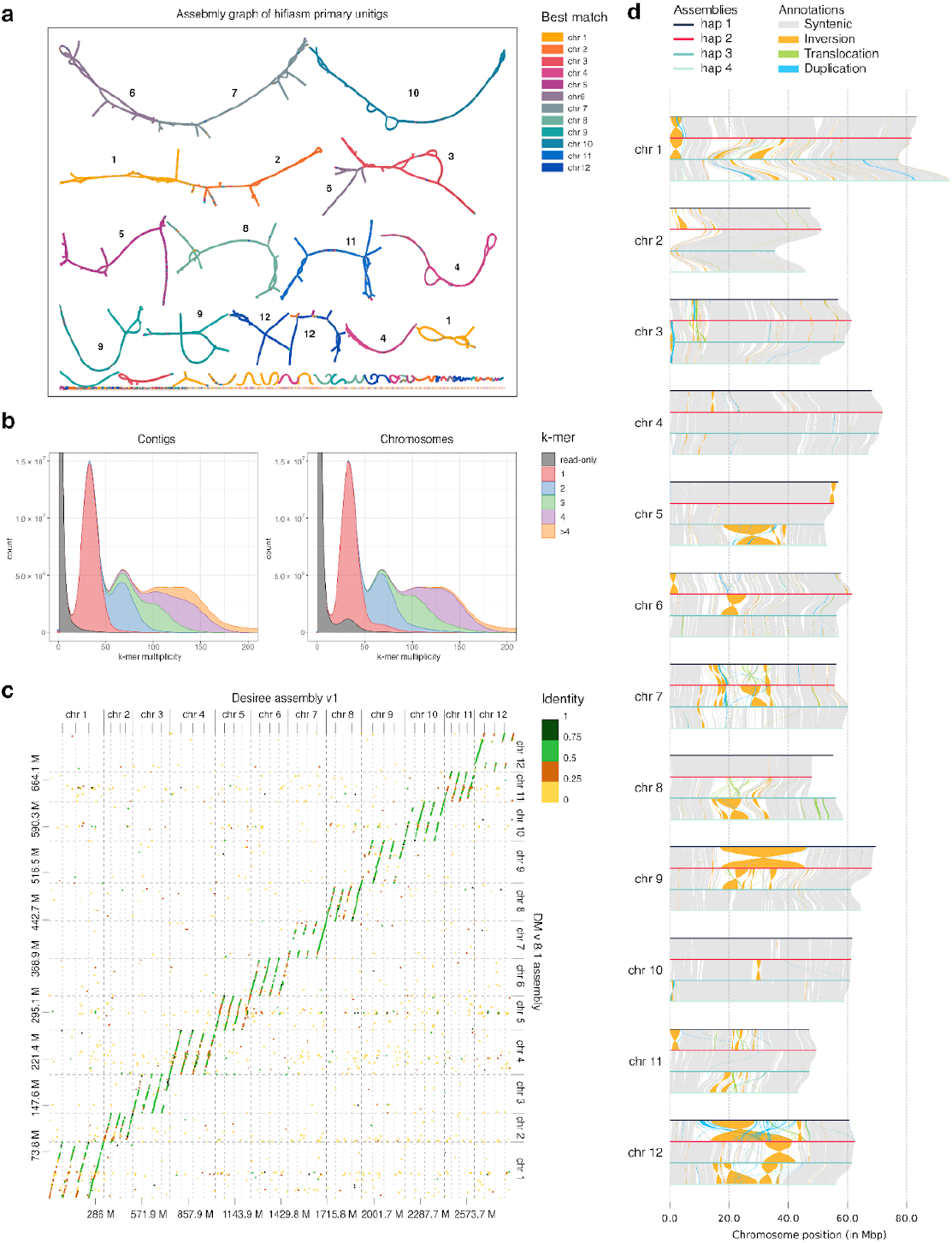
General characteristics of Désirée genome assembly **a)** Assembly graph of primary unitigs coloured by best match to DM chromosomes (also designated with numbers on the graph). **b)** Merqury k-mer spectra for initial contigs and scaffolded chromosomes. The k = 21 was used. K-mers are categorized as read-only (grey), unique (red), and shared (blue, green, purple, orange). Peaks corresponding to higher multiplicities indicate the presence of highly repeated k-mers. **c)** Dot plot comparing cv. Désirée chromosome-anchored contigs with DM v8.1 chromosomes. The colour designates contig identity. **d)** Genomic synteny of cv. Désirée haplotype-resolved assembly.

**Table 1.**
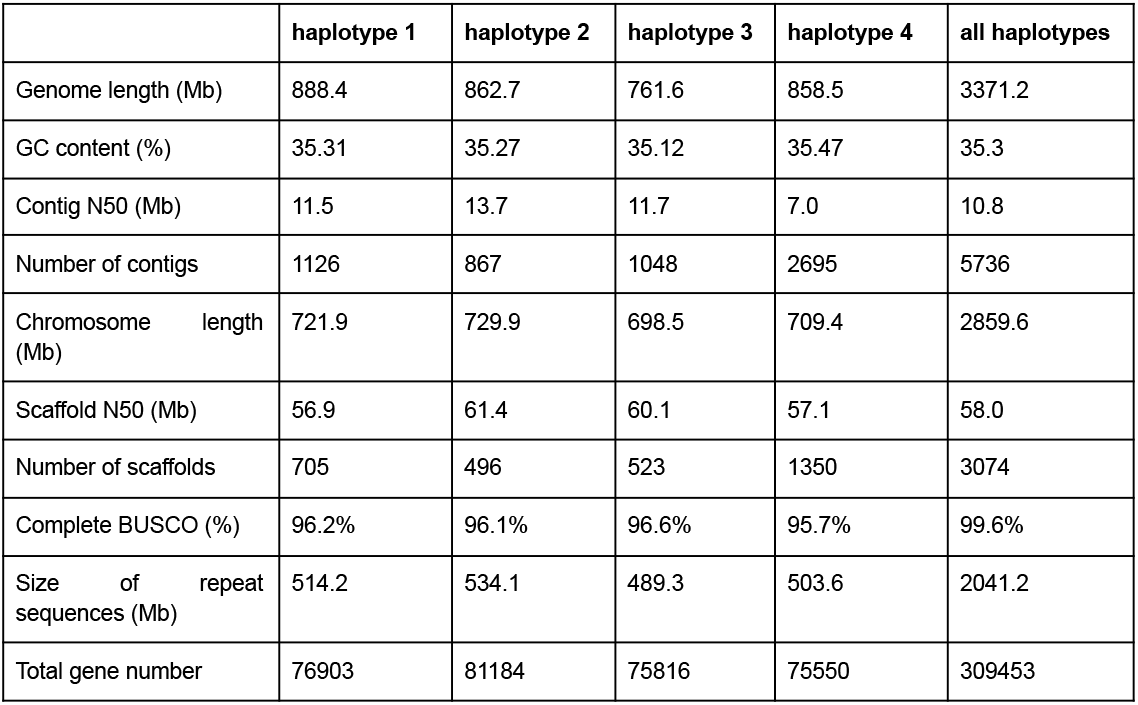
Summary of the four haplotypes of the Désirée genome assembly.

Contigs identified as contaminants were removed based on blastn (v0.8.1) searches against a custom-built contaminant database, which includes *Solanum* plastid and mitochondrial sequences and bacterial NCBI RefSeq sequences.

Decontaminated scaffolds were anchored to chromosomes by mapping Hi-C reads to each haplotype set separately following the manufacturer’s recommended pipeline for Omni-C data (https://omni-c.readthedocs.io). Briefly, Hi-C reads were mapped using BWA-MEM (v0.7.17-r1188)^24^ then the mappings were parsed with *pairtools* (v0.3.0)^25^ followed by samtools (v1.3.1)^26^ to identify and extract valid pairs. Valid pairs were used to anchor and orient scaffolds into chromosomes using YaHS (v1.2a.1)^27^ and Juicebox Assembly Tools (v2.17.00)^28,29^.

Chromosomes 11 and 12 of haplotype 4 lacked ∼20 Mbp and ∼30 Mbp part of the pericentromeric region, respectively, and haplotype 1 contained two additional unplaced scaffolds (scaffold_22 and scaffold_23). Alignment of these scaffolds to reference genome (DM v6.1) and inspection of Hi-C contacts suggested that these scaffolds are the missing regions of chromosomes 11 and 12 in haplotype 4. Therefore, we remapped Hi-C reads and incorporated these two scaffolds in haplotype 4 using Juicebox Assembly Tools (v2.17.00).

The final scaffolded assembly size amounts to 3.3 Gbp, with individual haplotypes ranging between 762 and 888 Mb. As expected, one haplotype is highly similar to the DM haplotype, whereas other haplotypes can be more dissimilar (Fig. 1c). A comparison of Merqury k-mer spectra between the initial contigs and the scaffolded chromosomes (Fig. 1a) reveals that many apparent duplications in the contigs are resolved during scaffolding. A small proportion of sequences remains missing from the chromosomes and those can be found in the whole genome FASTA.

The haplotype assemblies were sequentially aligned using minimap2 (v2.28) and analyzed with SyRi (1.7.0) to identify syntenic regions and structural rearrangements which were visualized using plotsr (v1.1.1, Fig. 1d).

### Genome annotation

Repeat elements in the *S. tuberosum* cv. Désirée genome were identified using the Extensive *de novo* TE Annotator (EDTA, v2.2.1)^30^. Repetitive sequences cover 489 - 534 Mbp per haplotype, representing more than 70% of the genome (Table 2).

**Table 2.** Summary of genome annotations for each haplotype.

| Type | haplotype 1 | haplotype 2 | haplotype 3 | haplotype 4 |
| --- | --- | --- | --- | --- |
| **Repeat elements** |  |  |  |  |
| DNA | 46.6 Mbp (6.4%) | 54.1 Mbp (7.4%) | 44.6 Mbp (6.4%) | 45.9 Mbp (6.5%) |
| Helitron | 36.1 Mbp (5.0%) | 38.0 Mbp (5.2%) | 33.6 Mbp (4.8%) | 42.3 Mbp (6.0%) |
| LINE | 12.4 Mbp (1.7%) | 8.1 Mbp (1.1%) | 7.5 Mbp (1.1%) | 8.1 Mbp (1.1%) |
| LTR | 176.5 Mbp (24.4%) | 188.0 Mbp (25.8%) | 165.9 Mbp (23.7%) | 193.6 Mbp (27.3%) |
| LTR/Copia | 16.8 Mbp (2.3%) | 19.3 Mbp (2.6%) | 20.3 Mbp (2.9%) | 23.9 Mbp (3.4%) |
| LTR/Gypsy | 136.2 Mbp (18.9%) | 133.6 Mbp (18.3%) | 130.0 Mbp (18.6%) | 102.8 Mbp (14.5%) |
| MITE | 11.9 Mbp (1.6%) | 10.2 Mbp (1.4%) | 13.0 Mbp (1.9%) | 10.6 Mbp (1.5%) |
| Other | 72.8 Mbp (10.1%) | 76.2 Mbp (10.4%) | 69.7 Mbp (10.0%) | 71.3 Mbp (10.1%) |
| SINE | 5.1 Mbp (0.7%) | 6.6 Mbp (0.9%) | 4.7 Mbp (0.7%) | 4.9 Mbp (0.7%) |
| Total | 514.2 Mbp (71.2%) | 534.1 Mbp (73.2%) | 489.3 Mbp (70.1%) | 503.6 Mbp (71.0%) |
| **Protein-coding genes** |  |  |  |  |
| Total gene number | 76903 | 81184 | 75816 | 75550 |
| Mean gene length (bp) | 1695.85 | 1610.97 | 1687.71 | 1677.79 |
| Mean CDS length (bp) | 1062.59 | 1032.74 | 1060.23 | 1061.68 |
| Mean exon number | 5.28 | 5.04 | 5.31 | 5.28 |
| Mean intron number | 4.28 | 4.04 | 4.31 | 4.28 |
| Complete BUSCO (%) | 94.1% | 93.3% | 95.4% | 93.7% |
| Single Omark HOGs | 82.9% | 82.5% | 84.3% | 82.8% |
| Duplicated Omark HOGs | 11.6% | 11.6% | 11.5% | 11.9% |
| Missing Omark HOGs | 5.5% | 5.9% | 4.2% | 5.4% |
| Mercator4 proteins annotated (%) | 93.5% | 93.5% | 93.7% | 93.5% |
| Mercator4 proteins classified (%) | 50.5% | 46.5% | 50.7% | 50.0% |
| Mercator4 bins occupied (%) | 94.2% | 93.9% | 94.6% | 94.3% |

The prediction of protein-coding genes in the assembled *S. tuberosum* cv. Désirée was determined using five complementary approaches: *de novo*, homology-based, transcriptome-based, deep-learning, and reference-based predictions (Fig. 2).

**Fig. 2.**
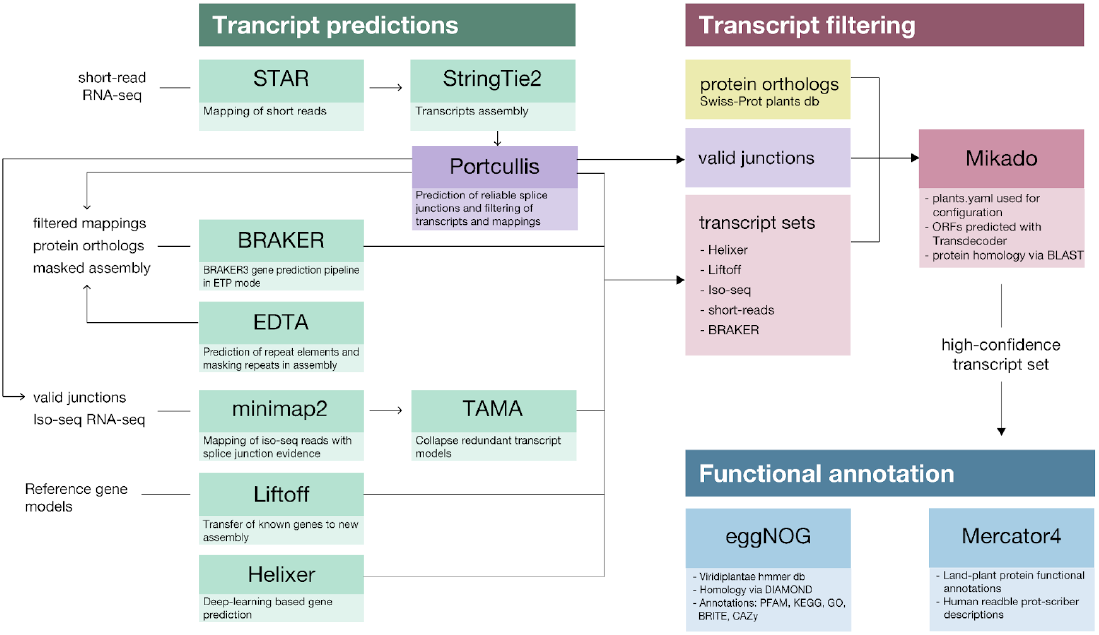
Workflow overview of *S. tuberosum* cv. Désirée genome annotation.

For transcriptome-based prediction, two methods were applied for short reads and Iso-Seq reads, respectively. Short reads from multiple tissues were aligned to each haplotype using STAR (2.7.10a)^31^, and transcripts were assembled with StringTie2 (v2.2.1)^32^, followed by Portcullis (v1.2.4)^33^ for junction validation. Iso-Seq reads from five *S. tuberosum* cultivars were mapped to both haplotypes using minimap2 (v2.28)^34^, and transcripts were generated using IsoQuant (v3.3.1)^35^ and TAMA Collapse (tc_version_date_2023_03_28) ^36^.

BRAKER3 (v3.0.8)^37^ was used in ETP mode to predict gene models by integrating *de novo*, homology-based, and transcriptome-based predictions. Repeat masking of the assembly was performed with RepeatMasker (v4.1.2), using EDTA annotations. Protein sequences from OrthoDB (green plant orthologs) were provided as evidence, and short-read STAR alignments with invalid junctions removed were included.

Helixer (v0.3.3)^38,39^ was used for deep-learning-based gene prediction via its web interface (https://www.plabipd.de/helixer_main.html). Gene models from the *S. tuberosum* reference genome (DM v6.1, UniTato annotation) were transferred to the Désirée assembly using Liftoff (v1.6.3)^40^. All five transcript or gene model sets were consolidated using Mikado (v2.3.4)^41^ to generate a non-redundant set of transcripts. Protein-coding gene completeness was assessed using BUSCO (Table 2, v5.4.7, solanales_odb10 dataset) and OMArk (v0.3.0, omamer v2.0.2)^42^.

The predicted protein-coding genes were functionally annotated using EggNOG Mapper (v2.1.11)^43^ with the EggNOG database (version 5.0.2)^44^ for the Viridiplantae subset. This included categories such as gene names, Gene Ontologies (GOs), enzyme functions (EC), and KEGG pathways, reactions, and modules, along with CAZy families, PFAM domains, and more. Additionally, functional land-plant protein annotations were predicted using Mercator4 (v7)^45^ via the web platform (https://www.plabipd.de/mercator_main.html). Annotations from EggNOG and Mercator4 were combined into the final GFF3 annotation file.

Orthologous groups between haplotypes and UniTato genes were identified using OrthoFinder (v2.5.5)^46^. Across haplotypes, 55.3% of orthogroups contained genes from all four haplotypes, 22.9% from three haplotypes, 19.2% from two haplotypes, and 2.7% from a single haplotype. When comparing the Désirée annotation to UniTato, 17.24% of genes were specific to the Désirée annotation.

## Data Records

The raw sequencing data, including Illumina Hi-C, Illumina paired-end, PacBio HiFi, and ONT reads, have been deposited at the National Center for Biotechnology Information (NCBI) Sequence Read Archive (SRA) under BioProject number PRJNA1185028. Plastid, mitochondrial and bacterial sequences used for removal of contaminant contigs were downloaded from NCBI RefSeq release 218. Transcriptomic data used for gene annotation was downloaded from public repositories: SRA under accessions PRJNA1192223, PRJNA1186376, PRJNA718240, PRJNA803222, PRJNA1209787 and PRJNA1191209; the Gene Expression Omnibus (GEO) under accession GSE232028; and the National Genomics Data Center (NGDC) under accession CRA006012. Existing gene models used in the gene annotation pipeline were downloaded from https://unitato.nib.si and https://spuddb.uga.edu. The genome assemblies of the four haplotypes have been submitted to NCBI GenBank under the BioProject accessions PRJNA1196677, PRJNA1196678, PRJNA1196679 and PRJNA1196680. The assembled genome, including annotations, methylation profile and identified orthologs, is hosted in a Zenodo repository under DOI: 10.5281/zenodo.14609304 and is also accessible via an interactive genome browser at https://desiree.nib.si.

## Technical Validation

We assessed the assembly quality and completeness using DNA sequencing read mapping, CRAQ, BUSCO analysis, and Merqury k-mer based evaluation. Illumina reads were mapped with BWA (v0.7.17), while PacBio and ONT reads were aligned using minimap2 (v2.28). Mapping rates were 99.90%, 100.00%, and 99.74% for Illumina paired-end, PacBio, and ONT reads, respectively. CRAQ (v1.0.9)^47^ analysis of PacBio and Illumina mappings yielded a regional AQI of 96.3 and an overall AQI of 97.5, classifying the assembly as reference quality (AQI > 90). Assembly completeness was assessed with BUSCO (v5.4.7) using the solanales_odb10 lineage database, identifying 5930 (99.6%) of the 5950 BUSCO orthologous groups in both the whole genome and chromosome-only assemblies (Table 1). Merqury (v1.3) analysis, using a Meryl (v1.3) database constructed from Illumina reads, estimated genome completeness at 98.57% for the whole genome and 95.73% for the chromosomes. The estimated QV values were 54.30 and 58.53 for the whole genome and chromosomes, respectively.

Completeness of gene annotation was assessed using OMArk (v0.3.0, omamer v2.0.2), BUSCO (v5.4.7) and Mercator4 (v7). OMArk analysis demonstrated that our annotation captured 94.1%-94.6% of Hierarchical Orthologous Groups (HOGs) per haplotype, with duplication rates ranging from 11.5% to 11.9% (Fig. 3a). When combining genes from all haplotypes, the proportion of complete HOGs reaches 99.3%, meaning that not all conserved genes are present in all haplotypes. Similarly, BUSCO analysis reported a haplotype completeness range of 93.3%–95.4% (Table 2), while the whole genome annotation achieved 99.4% completeness. Protein classification via Mercator4 revealed that 93.9%–94.6% of Mercator bins were occupied per haplotype, increasing to 97.5% when combining all proteins (Table 2). As expected, the Mercator bin with the largest proportion of missing proteins was associated with clade-specific metabolism (Fig. 3b). Additionally, the classified proteins showed no significant deviation from the median protein length, confirming consistency in annotation quality (Fig. 3c).

**Fig. 3.**
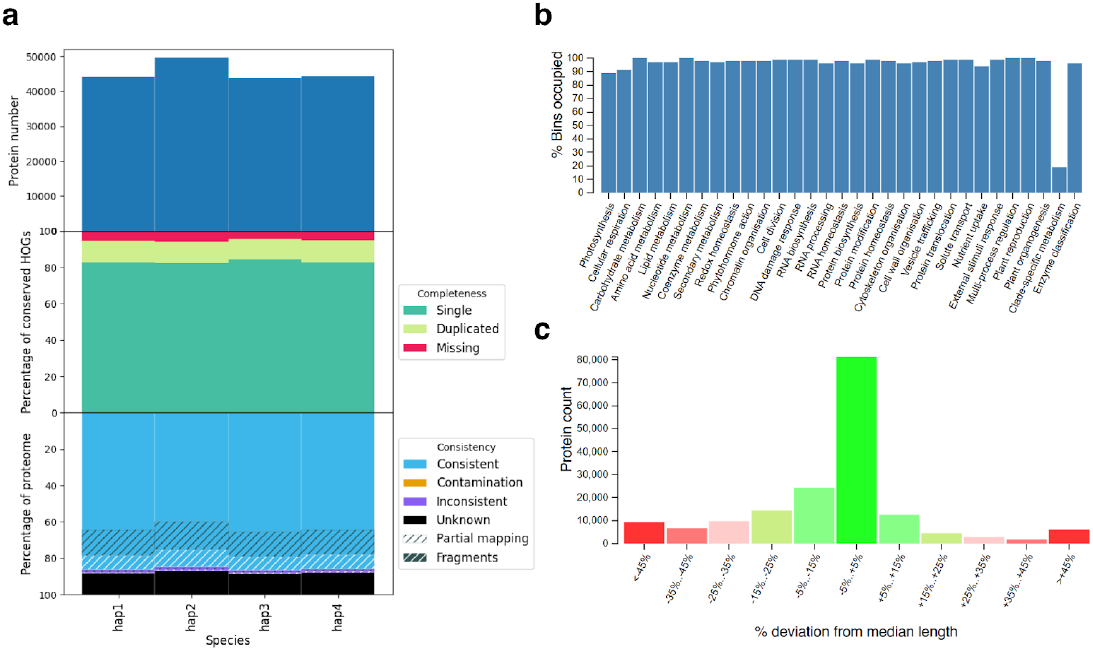
Validation of gene annotation. **a)** OMArk quality assessment showing consistency, completeness and count of proteins across all four haplotypes. **b)** Histogram showing the percentage of Mercator4 functional bins occupied by the Désirée proteins. **c)** Histogram displaying the distribution of proteins grouped by their percentage deviation from the median protein length.

## Usage Notes

The presented Désirée genome assembly is of high contiguity, completeness and phasing quality and presents a valuable resource for haplotype-aware transcriptomics, proteomics and epigenomics analyses. The transfer of UniTato annotations^48^ provides translation of gene identifiers from the DM to the Désirée genome. The RNA-seq datasets used to supplement gene model annotation are predominantly from mature leaf and root tissue, thus genes specifically expressed in other tissue and developmental stages may not be fully captured in the current annotation.

The genome was produced from a plant propagated in tissue culture for over a decade. A recent pangenome study^49^ found that *in vitro* propagated plants of the *Solanum* section Petota have greater numbers of TEs in their genomes. While this seems to hold for LTR elements and DNA transposons in the Désirée genome, overall TE expansion is not evident. Examining the DNA methylation profile available in the Désirée genome browser might provide more insight into specific transposable element expansion in this cultivar.

Recently, efforts were made to generate potato pangenomes^9,49^. However, the number of included phased tetraploid genomes is still limited. Including Désirée and more phased tetraploid genomes will improve the completeness of potato pangenome. This will bridge knowledge gaps in potato genomics and give potato breeders a powerful toolkit for developing more resilient and productive cultivars.

## Code Availability

The code, scripts and command-line tool commands used for genome assembly, annotation and quality control are freely available in the GitHub repository https://github.com/NIB-SI/desiree-genome.

## Acknowledgement

This work benefits from resources and services provided by ELIXIR, a distributed infrastructure for life science data, funded by national governments and the European Commission, particularly the Elixir-SI node for performing Illumina paired-end sequencing.

Funding for this work was provided by the European Union’s Horizon 2020 research and innovation programme project ADAPT (grant agreement No GA 2020 862-858), Slovenian Research and Innovation Agency (ARIS) project grants P4-0165, P4-0431, and J4-3089. SB and BU are supported by the German Federal Ministry of Education and Research (BMBF) in the frame of the German Network for Bioinformatics Infrastructure (de.NBI).

## Author contributions

**TG**: Methodology, Data curation, Investigation, Visualization, Writing - Original Draft. **SB**: Investigation, Writing - Review & Editing. **BU**: Writing - Review & Editing. **NYRG**: Resources, Writing - Review & Editing. **RS**: Resources, Writing - Review & Editing. **LA**: Resources, Writing - Review & Editing. **MT**: Funding acquisition, Writing - Review & Editing. **KG**: Funding acquisition, Conceptualization, Writing - Review & Editing. **MP**: Conceptualization, Validation, Resources, Supervision, Project administration, Writing - Review & Editing.

